# A gene-level test for directional selection on gene expression

**DOI:** 10.1101/2022.11.03.515083

**Authors:** Laura L. Colbran, Fabian C. Ramos-Almodovar, Iain Mathieson

## Abstract

Most variants identified in human genome-wide association studies and scans for selection are non-coding. Interpretation of these variants’ effects and understanding of the way in which they contribute to phenotypic variation and adaptation in human populations is therefore limited by our understanding of gene regulation and by the difficulty in confidently linking non-coding variants to genes. To overcome this, we developed a gene-by-gene test for population-specific selection based on combinations of regulatory variants.

We extended the *Q_X_* test for polygenic selection to test for selection on regulatory variants for 17,388 protein-coding genes across 2,504 individuals. We identified 45 genes with significant evidence (FDR <0.1) for selection, including *FADS1*, *KHK*, *SULT1A2*, *ITGAM*, and genes in the HLA region. We further confirm that significant selection signals do correspond to plausible population-level differences in predicted expression. However, we find that very few (0.2%) genes have strong evidence for directional, population-specific selection on the component of their expression that is predicted by *cis*-regulatory variants. While this is consistent with most *cis*-regulatory variation evolving under genetic drift or stabilizing selection, it is also possible that any effects are smaller than we can detect, or that population-specific selection is driven by tissue-specific or *trans* effects.

Our gene-level *Q_X_* score is independent of other methods for detecting selection based on genomic variation, may therefore be useful when used in combination with more traditional selection tests to specifically identify selection on regulatory variation. Overall, our results demonstrate the utility of one approach to combining population-level information with functional data to understand the evolution of gene expression.

## Introduction

Natural selection is one process by which populations respond to their environment. Therefore, identifying phenotypes, genes and variants influenced by selection is an important aspect of understanding how organisms and populations adapt. In humans, there has been an increasing interest in identifying population-specific adaptations. This is usually in the hopes of better understanding the mechanisms underlying the phenotype (Crawford et al., 2017; Ilardo et al., 2018; Simonson et al., 2010) and using that information to improve human health. However, linking genomic signals of selection to specific phenotypes and evolutionary pressures remains challenging. It is believed that changes in gene expression underlie most recent evolution (Corradin et al., 2016; King and Wilson, 1975), and are therefore the most likely changes to underlie selection on complex traits. On the other hand, across species gene expression seems to largely be under stabilizing selection or evolving neutrally (Chen et al., 2018; Rohlfs et al., 2014; Signor and Nuzhdin, 2018). While there are some genes with evidence for directional selection on their expression (Blekhman et al., 2008), these effects are tissue-specific, and the extent to which selection plays a role on genome-wide gene regulation remains poorly characterized (Price et al., 2022).

One approach to testing for selection on complex traits is to start with an observed trait difference, then to test whether that difference is greater than expected compared to genetic difference (Whitlock,2008), and work backwards to understand the mechanism. The limitation of this approach is that it is difficult to account for the environmental component of the phenotypic variance. Another approach, typical for genome-wide scans for selection, is to identify individual outlier haplotypes based on allele frequency or linkage disequilibrium (LD) patterns, then work forward to understand which variant is the causal one, and what it might be influencing (Voight et al., 2006). Since variants rarely act in isolation, many traits are polygenic and any signals of selection on complex traits could therefore be spread across many variants across the genome. This can be identified by looking for enrichment of these locus-specific selection signals (Field et al., 2016). Somewhat intermediate to these approaches, the *Q_X_* statistic (Berg and Coop, 2014) tests for polygenic selection by using genome-wide association results to test for systematically divergent allele frequencies among all independent variants associated with a phenotype, in theory capturing only genetic contributions to the phenotypic variance. However, in practice even this approach can be confounded due to poorly-controlled population stratification in the underlying GWAS (Berg et al., 2019). It is therefore unclear to what extent polygenic selection is relevant for human adaptation.

Most variants associated with complex traits through genome-wide association studies (and therefore those most likely to be subject to selection) are non-coding and likely operate through changes in gene expression. Therefore directional polygenic selection on complex traits, if it exists, may involve directional selection on gene expression. We aim to test for selection on the expression of specific genes, reasoning that this phenotype might reflect the effects of natural selection more clearly than other complex traits. However, gene regulation is extremely complicated, and mapping particular variants to their effects and genes is challenging Benton et al. (2019); Gasperini et al. (2020). In parallel to the development of GWAS methodology, there has also been a proliferation of methods and data to associate variants with gene expression. Single-variant eQTL studies are common, however genes often have multiple eQTL acting in concert to modulate expression. In addition, it is often prohibitively expensive to obtain the RNA-seq data needed to study expression directly in very large samples. Joint-tissue Imputation (JTI) is a machine-learning method that was developed to fill that gap by using expression and functional genomics data across dozens of tissues to predict gene expression based on combinations of genetic variants (Zhou et al., 2020a). These models and similar ones can be used in a transcriptome-wide association study (TWAS) to identify gene-level associations with complex traits (Petty et al., 2019; Zhu and Zhou, 2020), and we have used them to study predicted differences between ancient populations (Colbran et al., 2021). However whether predicted differences between populations in fact reflect real differences in expression and if so whether they are the result of directional selection are still open questions.

The goal of our study is to use the *Q_X_* test with eQTL instead of GWAS data to test for population-specific directional selection on combinations of variants that are associated with expression of specific genes. Done genome-wide, this results in a gene-level test for selection on regulatory variation, which we believe will be more specific than polygenic selection scans on higher-level traits and more interpretable than variant-level scans. Overall, this work identifies dozens of genes whose regulation has been influenced by population-specific selection, and demonstrates the utility of incorporating mechanistic data into genome-wide scans for selection.

## Results

### Testing for Selection on Regulatory Variants

The first step in characterizing selection on gene regulation is to identify the relevant genetic variants to include. To do this, we used published statistical models of gene regulation built using Joint Tissue Imputation (JTI) (Zhou et al., 2020a). The training process included variant selection and quantification of independent linear effects of combinations of variants on expression of each gene in each tissue. We used models that were trained using the genotypes and transcriptomes from 49 tissues from the GTEx project (Aguet et al., 2017). Because these models borrow information across tissues, they are often correlated with each other, particularly for genes with shared regulatory patterns. We wanted to use one model per gene in order to limit the multiple-testing burden, since expression patterns across tissues are not independent. While ideally we’d like to study the biologically relevant tissue for each gene, in many cases the tissue to choose is not obvious. We therefore decided to focus on models that capture the most accurate information about individual-level gene expression. We compared predicted expression to observed expression in Lymphoblastoid Cell Lines (LCLs) for 5 populations from 1000 Genomes (1kG; N = 447) (Lappalainen et al., 2013). We found that the rank correlation between predicted and actual expression for the gene models trained in GTEx LCLs was highly variable by gene (Fig. 1A; median Spearman *ρ* = 0.12, maximum *ρ* = 0.93). It was also significantly correlated with model performance during training (Fig. 1B; *ρ* with model *R*^2^ = 0.58, *P* = 2.1 × 10 ^-686^), indicating that the training *R*^2^ is a useful proxy for out-of-sample performance in tissues we have not measured directly. We decided to use the model for each gene with the highest training *R*^2^, regardless of the primary tissue it was trained in (the “best models”). While we focused on these best models for our genome-wide scan for selection, we suggest focusing on relevant tissues when testing specific hypotheses.

**Figure 1:**
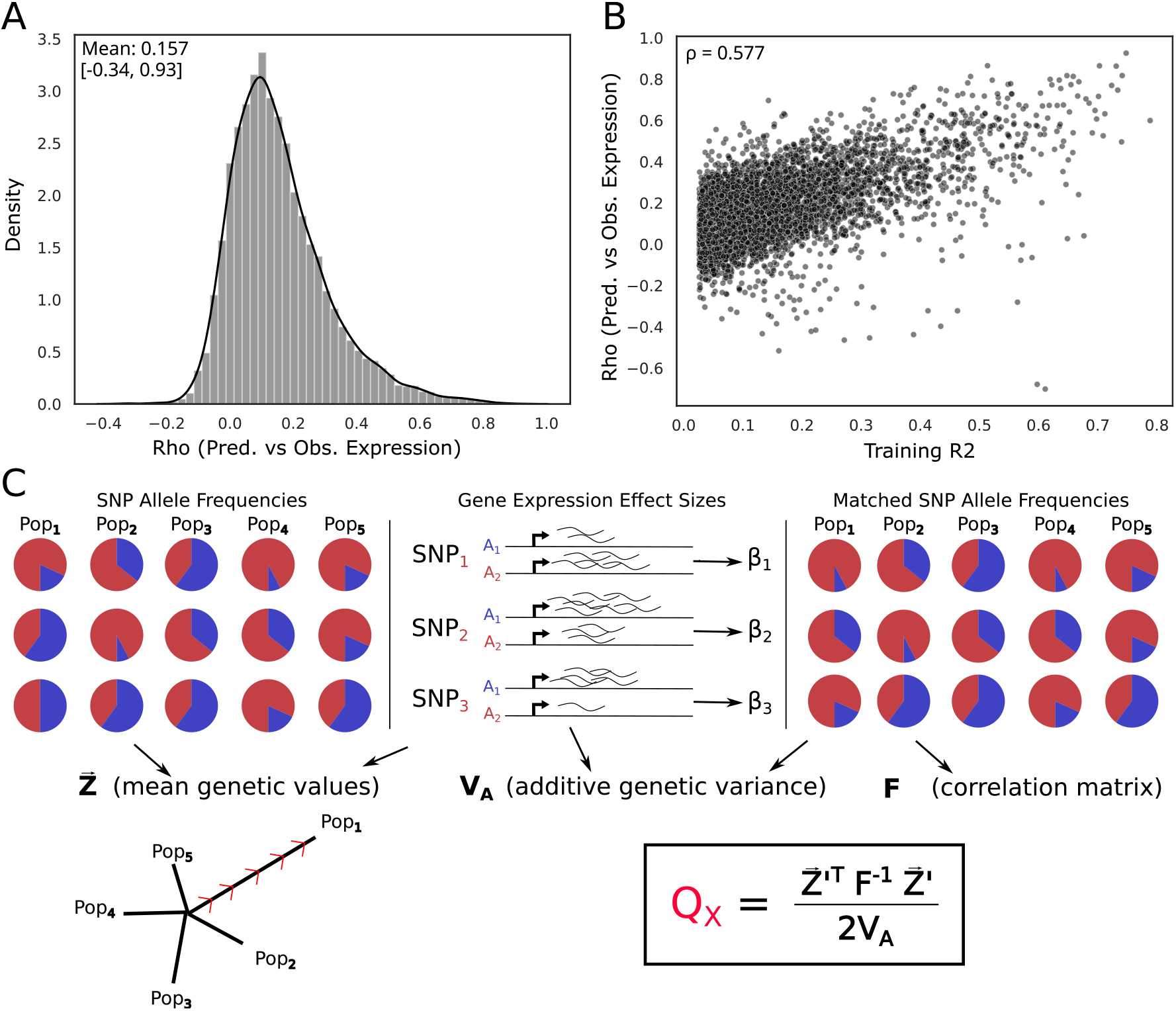
We adapted the Qx statistic to test for selection on regulatory variants. A) Spearman *ρ* between observed and predicted expression in 1kG for 7,251 JTI models trained in GTEx LCLs. B) *ρ* between observed and predicted expression in 1kG for the best JTI models for 26,878 genes C) Schematic of Qx calculation as applied to PrediXcan models.

To test for selection on gene regulation, we adapted the *Q_X_* test for polygenic selection (Berg and Coop, 2014) (Fig. 1C). We calculated a per-gene *Q_X_* score for 2,504 individuals from 26 populations from the high coverage hg38 1kG data (Byrska-Bishop et al., 2022; The 1000 Genomes Project Consortium,2015) by using the effect sizes from the corresponding JTI model in place of the GWAS associations, and replicated our analyses in 929 individuals from 7 populations from the Human Genome Diversity Panel (Bergström et al., 2020). This allows us to test for overdispersion in genetic values of gene regulation, or coordinated differences in allele frequencies of regulatory variants (Methods). Theoretically the *Q_X_* statistic follows a *χ*^2^ distribution with degrees of freedom one fewer than the number of populations under consideration (Supp. Fig. 1). In practice however, it can be over- or under-dispersed for reasons other than selection, such as population stratification or stabilizing selection.

One way to control for this is to calculate an empirical distribution. In Berg and Coop (2014) this was done by resampling allele frequencies for the variants in question genome-wide (Supp. Fig. 3A). In our case, doing a similar process results in extremely badly calibrated *P*-values (Supp. Fig. 4); this is primarily because our gene regulation models break the assumption of independence between variants (discussed in more detail in Methods). While the effect sizes fit by the models are independent and variants were pruned for very high LD (*r*^2^ > 0.8), most models still contain variants with lower levels of LD. Randomizing allele frequencies does not account for these residual correlations and is therefore anti-conservative.

We therefore implemented two alternative strategies to compute P-values. First, instead of randomizing frequencies, we randomized effect sizes of the variants in each model by sampling from the distribution of effects across all models (Supp. Fig. 3B). This tests specifically for coordination in the effect sizes of the variants conditional on allele frequencies. Second, we calculated empirical *P*-values by fitting a gamma distribution to the *Q_X_* distribution.

To evaluate these two approaches, we used SLiM (Haller and Messer, 2022) to simulate varying degrees of population-specific selection for 20,000 genes with multiple regulatory variants, then calculated *Q_X_* and both gamma-corrected and effect-permuted *P*-values (Methods). We found that the gamma-corrected approach was uniformly more powerful than the permutation approach. Indeed, while the gamma-corrected test approaches a power of 1.0 under regimes with stronger selection, the effect-permuted version never reached that (Fig. 2A). On the other hand, we noted that the permutation approach was more robust to technical characteristics of the model such as the number of variants in a model and its training *R*^2^ (Fig. 5). In order to understand the difference in power we used the fact that the *Q_X_* statistic can be decomposed into two components. The *F_ST_*-like component captures allele frequency differences and the LD-like component incorporates the combinations of effect sizes and directions (Berg and Coop, 2014). Genes that are genome-wide significant (FDR<0.1) with gamma-corrected significant genes are outliers for both components. Genes that are significant with the permutation approach are not outliers in the *F_ST_* component and only slightly enriched in the LD component (Fig. 2B). In summary, the effect-permuted test seems to be very conservative, doesn’t capture high-Qx outliers and does not identify genes known to have strong signals for selection such as *FADS1*. We therefore focus in the main text on the results identified by the gamma-corrected test.

**Figure 2:**
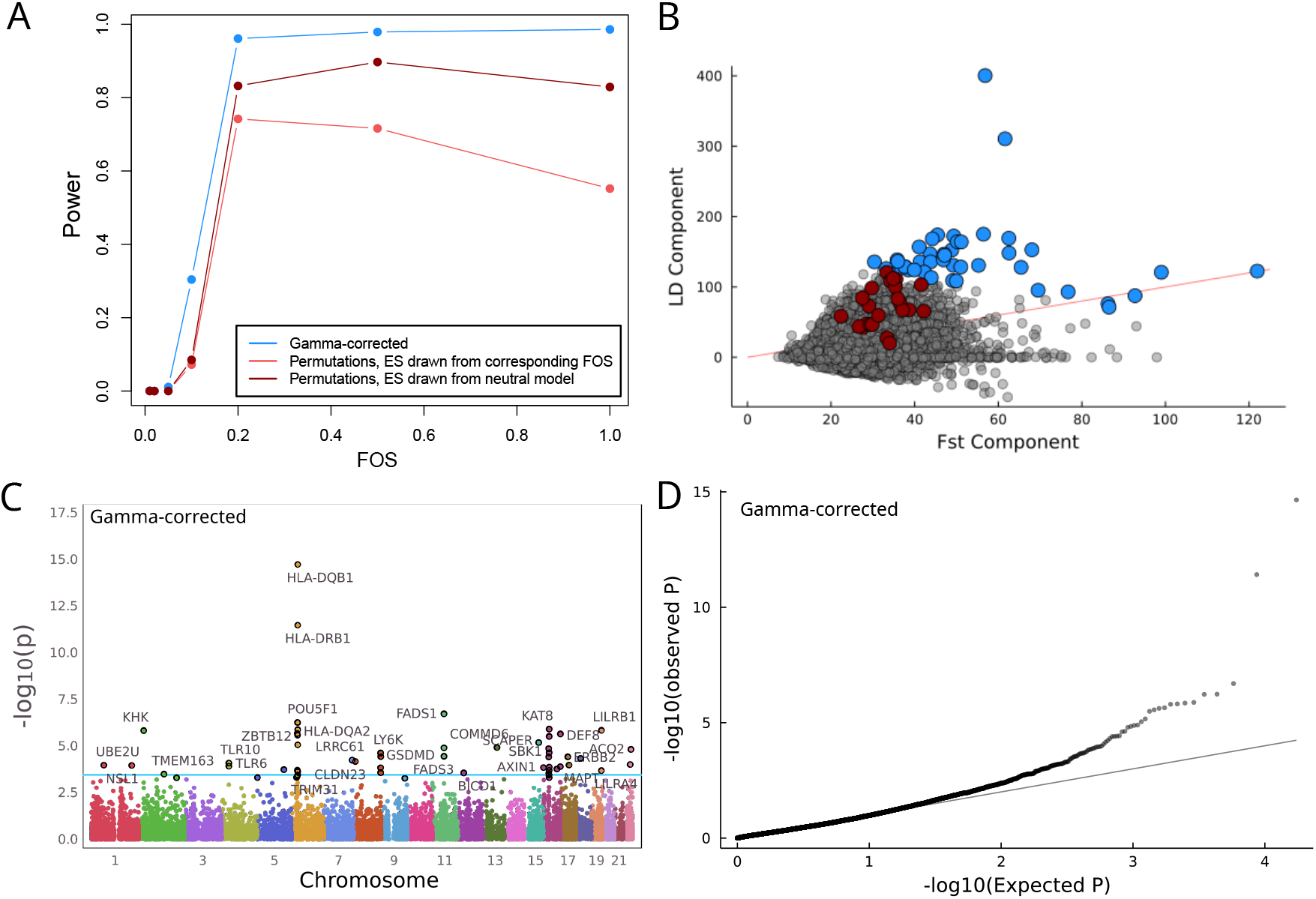
Genes under significant selection vary depending on *P*-value methodology. A) Manhattan plot and B) QQ-plot for gamma-corrected *Q_X_ P*-values per gene. Blue line corresponds to FDR < 0.1. C) The *Q_X_* score can be decomposed into its *F_ST_*-like component and its LD-like component. Significant (FDR < 0.1) genes in 1kG for the gamma-controlled and effect-permuted *P*-values are highlighted in blue and red, respectively, while the red line indicates where *F_ST_* = LD. The *Q_X_* score for each gene is obtained by adding the two components together. The Spearman rank correlation between the components is 0.15. D) Power curves for each *P*-value method, based on simulations (Methods). We calculated the power for the gamma-corrected version of the *Q_X_* test, as well as for 2 variations of the effect-permuted test. In the first, we drew the effect sizes from the simulation that modelled the corresponding selection strength, and for the other from the neutral effect model. FOS = fitness optimum shift.

With the gamma-corrected P-values we identified 45 genes with significant evidence of selection (FDR < 0.1; Fig. 2C-D). Because predicted expression of nearby genes shared regulatory haplotypes, these corresponded to 20 visible ‘peaks’ of nearby genes. These included several loci known to have experienced population-specific selection (e.g. *FADS1* and the TLR and HLA loci; Mathieson et al.,2015). These P-values are potentially still inflated by uncorrected population stratification, and are additionally correlated with both the number of variants in a model as well as its training *R*^2^ (Supp. Fig. 5). These technical aspects of the model training should not necessarily influence patterns of selection, but probably do affect power. These characteristics make it difficult to identify which gene in a peak is most likely to be the one directly under selective pressure, rather than merely influenced by the resulting allele frequency shifts. For comparison, when using the permutation test 23 genes have significant (FDR < 0.1) P-values. Overall the QQ plot shows little evidence of strong outliers (Supp Fig. 6). While there was no overlap in the significant genes in each P-value scheme, across all genes ordering was relatively highly correlated (Spearman *ρ* = 0.71), suggesting the two methods capture some of the same information.

We also ran the *Q_X_* scan in HGDP in order to identify whether selection patterns and particular genes replicated in an independent dataset. For the gamma-corrected *P*-values, 48 genes were significant at FDR < 0.1, and 4 peaks (HLA, *KHK, KAT8*, and *ACO2*) overlapped genes identified in 1kG (Supp. Fig. 7). For the effect size permuted P-values, no genes passed that significance threshold, though *SAMD10* (also identified in 1kG) did have the smallest P-value. Overall these results suggest that while some genes have experienced directional selection on expression driven by the *cis*-regulatory variation, it is relatively rare.

### *Q_X_* is independent of other selection metrics

While our *Q_X_* analysis did identify several previously-known genes, we wanted to know whether including gene regulation information was generally giving us more information than other, less-specific tests for selection. We therefore calculated the correlation between gene-level *Q_X_* and a variety of other measures of selection (Fig. 3). We find that the gamma-corrected P-values are not strongly correlated with either Loss-of-Function intolerance or PhyloP, both of which are metrics of evolutionary constraint (Lek et al., 2016; Pollard et al., 2010), or with two haplotype-based tests for more recent selection (iHS, nSL, averaged across a window for each gene; highest Spearman *ρ* = −0.032 between *P* and PhyloP), although as expected these other four metrics do show some correlations with each other (highest Spearman *ρ* = 0.38 between iHS and nSL). The same is true of the effect size permuted *P*-values.

**Figure 3:**
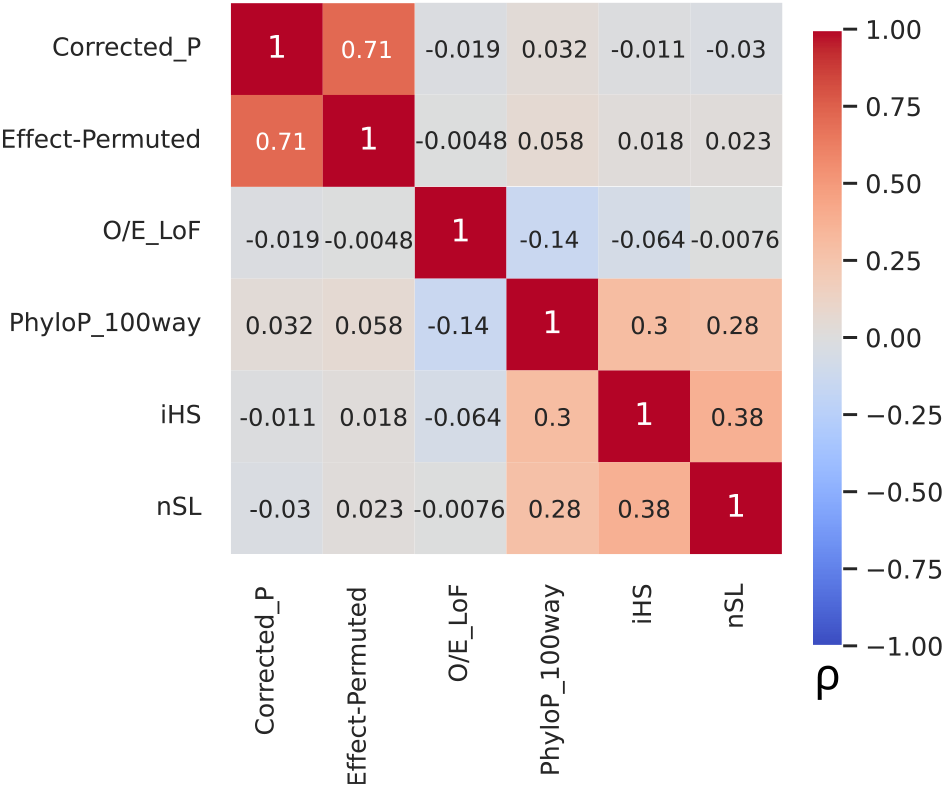
The *Q_X_* statistic is not correlated with other selection statistics. Pairwise heatmap of Spearman rank correlations between *Q_X_ P*-values and various selection-related scores.

We also tested whether different classes of genes were enriched for signals of directional regulatory selection. LoF-intolerant genes are somewhat depleted among the genes with the smallest gamma-corrected *P*-values (e.g. OR = 0.281, *P* = 0.0011 for the top 100 genes; Supp. Fig. 8), suggesting that genes under strong constraint on their protein sequence also tend to be more constrained on their regulatory variation. Surprisingly, housekeeping genes, a broadly-expressed class of genes responsible for basic cellular functions that we might expect to be similarly constrained, are somewhat enriched among genes with more evidence for selection (OR = 2.64, *P* = 2.8 × 10^-4^ for top 500 gamma-corrected genes; OR = 5.92, *P* = 1.5 × 10^-3^ for top 2-effect-permuted genes). This is consistent with patterns seen in our previous study of regulatory differences between ancient populations (Colbran et al., 2021). We also tested for enrichment of genes of certain functional categories, such as viral-interacting proteins (Enard et al., 2016), immune genes that respond to interferon, and diet-related genes with known selection signals (Rees et al., 2020), as well as genes that have undergone stabilizing selection on expression across species (Chen et al., 2018), but found no significant trends for any of these categories for either set of P-values. Technically, the selected diet genes were significantly enriched among the top 10 gamma-corrected genes (*P* = 0.021); however that signal is driven entirely by *FADS1*, and is therefore uninformative about broader patterns.

### Patterns of predicted expression among selected genes

We next wanted to confirm whether significant *Q_X_* scores corresponded to differences in predicted expression between populations; this is expected because the *Q_X_* scores are built on the JTI models. To test this, we applied the best JTI models to predict expression in the same individuals we used in calculating the *Q_X_* scores and summarized these predictions across populations. Indeed, we found that the genes showing significant selection differed between populations (Fig. 4A). For example, *LY6K* has a median predicted expression 3 standard deviations higher in Japanese populations than in most others. This is true for genes identified as significant under both *P*-value schemes (Supp. Fig. 9, suggesting both methods identify genes with overall predicted differences. These predicted patterns are also largely similar in HGDP, despite the decreased resolution of that data (Supp. Fig. 10). As expected, these predicted differences do not always agree with patterns of observed expression in LCLs (*P* = 0.241 for 1kG); this could be because of differing tissue expression patterns (we only have observed expression in LCLs), or because of differing environment or genetic background compared to the model training data (Supp. Fig. 11).

**Figure 4:**
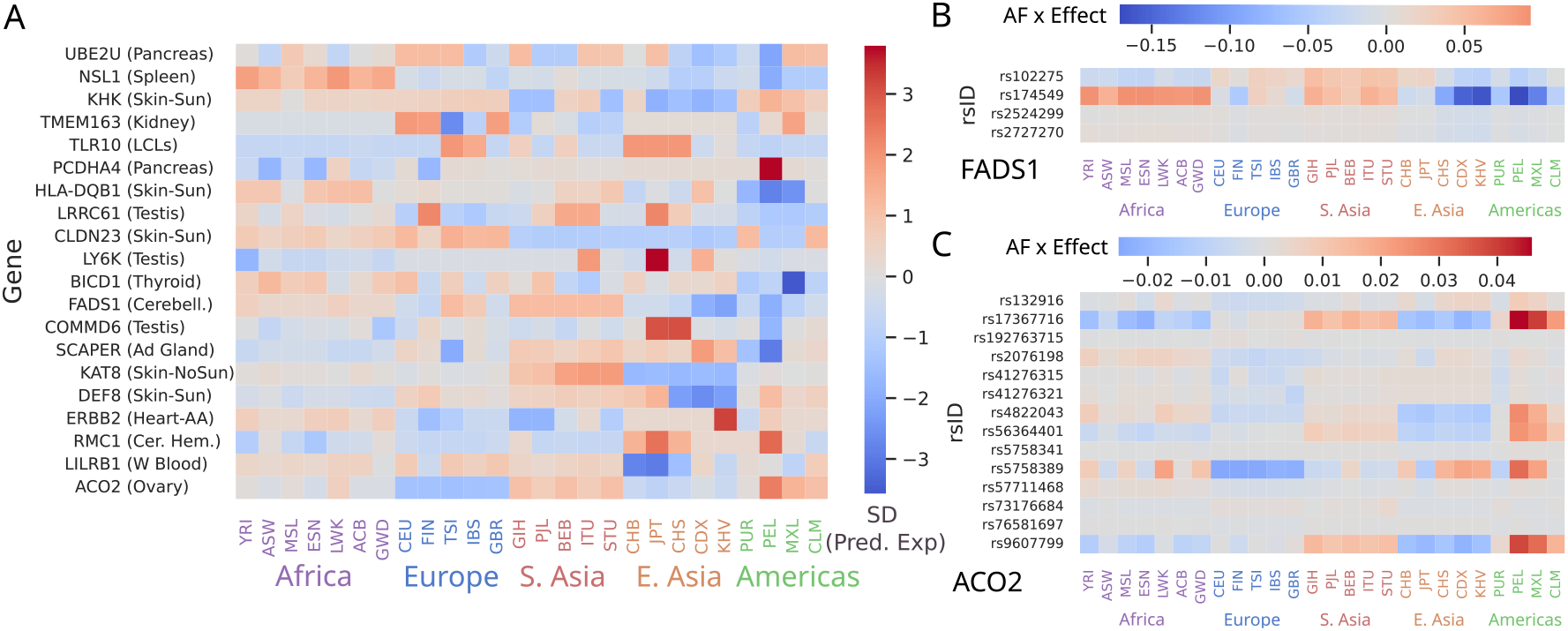
Different combinations of variant effects can drive predicted differences. A) Median predicted expression in each 1kG population for the top gene in each peak of the gammacorrected *P*-values. For display purposes, for each gene is standardized across populations. B) for *FADS1* there is one primary haplotype (tagged by rs174549), while for C) *ACO2* there are 3 variants driving the upregulation in PEL. Cells are coloured by the product of JTI effect size times effect allele frequency in each population. Values in B) and C) are mean-centered for each variant.

While there are 20 peaks of significant selection in the gamma-corrected *P*-values calculated in 1kG, it is unlikely that every gene in each peak is actually under selection. Instead, it is likely that the regulation of one gene has an impact on fitness, and the expression of other genes with shared or linked regulatory variation hitchhikes along with the selected gene. For example, *FADS1* is a well-established example of a selected haplotype whose effect is correctly modeled and detected (tagged by rs174549; Fig. 4B). However the nearby genes *FEN1* and *FADS3* are significant in this analysis as well, as they are also influenced by the selected haplotype. Additional lines of evidence are therefore necessary to understand which gene in a peak is the cause of the selection, and which are side-effects. We focused on the 4 peaks that replicate in HGDP (the HLA locus, *ACO2, KHK*, and *KAT8*). The HLA region is another well-established locus, but the other three are novel. *ACO2* and *KHK* are both the sole genes in their peaks that replicate, so are the most likely candidates in each. *ACO2* is a mitochondrial gene that plays an important role in the TCA cycle (Gruer et al., 1997), and is predicted to be relatively downregulated in Europeans, and upregulated in Peruvians from Lima (PEL) and other Native American populations. Unlike for *FADS1*, these predictions are result of multiple SNP effects, although dominated by the allele frequencies of rs5758389 (Fig. 4C). *KHK* is a gene responsible for catabolizing dietary fructose (Bonthron et al., 1994), and is predicted to be downregulated in Asian populations (JPT, CHS, CDX, KHV, GIH, PJL, STU). While each of these genes was relatively isolated, the *KAT8* peak had 7 genes that replicated between 1kG and HGDP *(KAT8, ZNF668, ITGAM, STX1B, SNF646, SBK1*, and *SULT1A2*). A closer look shows that this peak in fact represents two independent signals (Supp. Fig. 14), with *SBK1* and *SULT1A2* showing strong predicted differences in PEL, and the other 5 genes showing predicted differences among Asian populations. Each has at least one potential candidate for selection; *ITGAM* is an integrin that is part of the innate immune system (Ramírez-Bello et al., 2019), while *SULT1A2* is important for metabolizing hormones, drugs, and other xenobiotic compounds (Glatt et al., 2001). Both are involved in responding to the environment, and are therefore the most likely to be subject to population-specific selection.

In contrast, the effect size-permuted *P*-values did not show a tendency to form peaks. Only 23 genes were significant in 1kG, and none of these replicated in HGDP. These 23 significant genes perform a variety of different functions that are also potentially interesting in the context of population-specific selection, including the regulation of insulin secretion *(STXBP4*, the binding of HDL cholesterol *(HDLBP*), and viral replication (PPIE) (Wang et al., 2011). *SAMD10* is the gene that comes closest to replicating in 1kG and HGDP (*P* = 2.0 × 10^-6^ in 1kG, *P* = 1.2 × 10^-5^ in HGDP), and is a plasma membrane protein that is most highly expressed in the Cerebellum and in LCLs (Aguet et al., 2017). Compared to Europeans (the primary population used in training the JTI models), it is predicted to be upregulated in African and South Asian populations, particularly Gujarati, Indian Telugu and Sri Lankan Tamil, and downregulated in most East Asian populations (Supp. Fig. 9).

## Discussion

In this study, we applied the *Q_X_* test for polygenic selection to regulatory variants identified using JTI expression imputation models to test for population-specific selection on gene regulation in 26 human populations. We identified 45 genes with significant regulatory selection. These included loci such as *FADS1*, *TLR*, and the HLA region that have been previously identified, as well as novel loci such as *KHK*, *SULT1A2*, and *ITGAM*. It was common for nearby genes to share high *Q_X_* scores likely reflecting some combination of linkage disequilibrium and shared regulatory variants. We also used a more conservative approach, which uses only the magnitude and direction of effects (conditioning on allele frequency differences). This version only has power to detect genes with coordinated changes across multiple variants and found few genes with evidence of selection. Some of the exceptions include genes associated with metabolism *(HDLBP, STXBP4*) and immunity *(PPIE*). Despite correctly identifying some well-established examples, our gene-level *Q_X_* score is not highly correlated with other metrics of selection, suggesting that it captures independent information.

There are some caveats with this approach. First, we are limited to testing cis-eQTL identified in the predominantly European GTEx data used to train the JTI models. We therefore cannot test for selection on trans-regulatory effects, or on any population-specific eQTL that were rare or absent in GTEx. Our analysis is also potentially vulnerable to confounding due to population stratification in the gene expression data, which is difficult to correctly account for. Our gamma-controlled analysis is likely susceptible to similar problems seen in the original *Q_X_* studies (Berg et al., 2019). By resampling effect sizes, we controlled for those effects, but at the cost of much-reduced power. In addition, while a high *Q_X_* does correspond to population-level differences in predicted expression in a particular tissue, these differences are not always reflective of actual differences in gene expression. More work in diverse populations and environments will be needed to confirm which of our specific results are true changes in gene expression or merely regulatory turnover.

In addition, our gamma-corrected approach highlights the fact that nearby genes are often co-expressed and share regulatory regions (Delaneau et al., 2019), making it difficult to determine which gene is actively subject to selection. Calculating P-values by permuting the effect sizes does control for these possibilities, but it also severely limits the type of effects we have power to detect. While our approach does allow us to identify genes influenced by potentially selected regulatory haplotypes, this is analogous to the issues with overlapping eQTL and GWAS studies (reviewed by Cano-Gamez and Trynka, 2020). A combination of LD and shared regulatory structure means signals often encompass multiple genes, and the tissue- and context-specificity of gene regulation means that our study, while genome-wide, is not exhaustive. Further lines of evidence will be required to disentangle these associations.

Despite these caveats, we do confirm several known instances of selection. In the case of *FADS1*, our method correctly identifies the known regulatory haplotype, and correctly predicts the direction of differences in expression between populations (Ameur et al., 2012; Buckley et al., 2017; Mathieson and Mathieson, 2018; Ye et al., 2017). The *TLR* locus is the site of a putative case of adaptive Neanderthal introgression (Quach et al., 2016), and our results suggest that this haplotype alters expression of all three genes in the locus. We identified several novel signals for genes involved in pathways that are likely to influence fitness in different environments. *KHK, ACO2, HDLBP, STXBP4*, and *SULT1A2* are all genes involved in various aspects of metabolism, whether directly diet-related or further downstream, while *PPIE* and *ITGAM* are both involved in innate immunity.

Overall our work suggests that strong, coordinated, population-specific selection on regulatory variation across multiple haplotypes is relatively rare among human genes and that patterns of variation in *cis*-regulation of gene expression across populations are largely explained by genetic drift. While it is possible that recent selection acts on regulatory variants we do not consider here (e.g. via trans effects), it is also possible that population-specific selection is not particularly common at strengths we are currently able to detect. Finally, our approach demonstrates that biologically-informed tests for selection can contribute orthogonal information to those based around LD patterns or other information, and therefore could be integrated with the results of other selection scans to increase interpretability.

## Methods

### Regulatory Variant Selection

To select regulatory variants and effect sizes, we started with the published Joint Tissue Imputation (JTI) gene expression models, which were trained in 49 tissues in GTEx v8 using all common variants (MAF > 0.05 in GTEx) (Zhou et al., 2020a). We used these models to predict expression in Lymphoblastoid Cell Lines (LCLs) for 447 individuals from 1000 Genomes Project (Geuvadis; Lappalainen et al., 2013;The 1000 Genomes Project Consortium, 2015). These individuals represented 4 populations of European ancestry (GBR, FIN, CEU, TSI) and one of African ancestry (YRI). We calculated TPM for ENA project PRJEB3366 using EMBL-EBI’s REST API (accessed March 8, 2022). We compared the predicted to observed expression for these individuals by calculating a Spearman rank correlation for 7,251 gene models trained in LCLs across all individuals. We compared the agreement of the models to the variance explained by the models in the training data by calculating the rank correlation across all genes between the model *R*^2^ and the predicted/observed correlation. For each gene, we selected the “best” model by choosing the model with the highest *R*^2^ across all tissues. This resulted in 26,878 genes for which we could compare observed and predicted expression.

### Qx score adaptation

We obtained effect sizes for variants by filtering the best models to include only protein coding genes (based on the “protein_coding” annotation in GenCode v26). The resulting 17,388 genes had a median of 12 (maximum 101) variants with effect sizes to calculate *Q_X_*. We calculated *Q_X_* as described by Berg and Coop (2014), using the effect sizes from each gene expression model in place of GWAS effect sizes. This allowed us to calculated a per-gene *Q_X_* score. For each gene, we chose 100 matched variants for each model variant by binning all variants into 25 bins based on alternate allele frequency in GTEx (i.e. a bin size of 0.02). We calculated *Q_X_* across the 26 populations of 1kG, encompassing 2504 unrelated individuals (Byrska-Bishop et al., 2022). We replicated using 929 individuals from the Human Genome Diversity Panel (HGDP) (Bergström et al., 2020), grouped into 7 broadly regional ‘populations’ due to the small sample size in individual populations. We plotted Manhattan plots with OpenMendel (Zhou et al., 2020b).

### P-value calculation

While *Q_X_* was designed to be a test for polygenic selection testing genome-wide, independent sets of variants, our adaptation of it would be better described as a multi-variant test for selection. The set of possible variants for each gene were pruned for linkage disequilibrium (LD) at *r*^2^ > 0.8 before the JTI models were trained (Zhou et al., 2020a), and models are built around independent, additive effects for the variants ultimately included. However, these variants are much closer together (within 2Mb), and models often include variants in moderate LD (*r*^2^ ≈ 0.4) with each other. This means that, while the effect sizes are independent, the allele frequencies are not necessarily, and the degree to which the frequencies are correlated with each other varies across genes. This makes calculating a P-value for gene-level *Q_X_* statistics complicated.

We tried three different methods for calculating *P*-values. The first is the method used in the original *Q_X_* study, wherein we construct a ‘null’ distribution of *Q_X_* scores for each gene by permuting the allele frequencies of the variants in the model (abbreviated as “freqPerm”; Supp. Fig. 3). For each permutation, we drew a random frequency for each variant in the model, holding effect sizes constant, and repeated that 100,000 times (up to 1,000,000 times if *P* < 10^-4^). As expected, because this permutation strategy breaks the LD structure present in many gene models, the resulting P-values are extremely poorly calibrated (Supp. Fig. 4).

We therefore calculated “corrected”*P*-values by instead fitting a gamma distribution to the *Q_X_* scores (with degrees of freedom equal to the number of populations minus 1). These *P*-values are much less inflated (Fig. 2B), while the ordering of genes is highly correlated with the order the freqPerm *P*-values gave (Spearman *ρ* = 0.993). They are however, somewhat correlated with technical characteristics of the gene models (Supp. Fig. 5).

To control for the technical confounding, we also calculated p-values by permuting the effect sizes for each gene while holding allele frequencies of variants constant (abbreviated as “effPerm”). Specifically, we randomly sampled effect sizes from the distribution of all possible effect sizes in any model, while holding the effect direction for each variant constant. We sampled 100,000 times for each gene, up to 1,000,000 for those with *P* < 10^-4^. While this did control for the technical variables (Supp. Fig. 5) and was still correlated with the corrected *P*-values (Spearman *ρ* = 0.709), it has the side effect of narrowing the hypothesis we were testing. Rather than a broad test for selection on regulatory variants, this permutation scheme emphasizes coordinated differences between populations (i.e. in the same effect direction) across multiple variants in a model. This means that true selection on a single regulatory haplotype (e.g. in the case of *FADS1*) is not identified.

### Power Calculations

We calculated the power of the gamma-corrected and effect-permuted test using simulations run in SLiM 4.0 (Haller and Messer, 2022). Simulations begin with an “ancestral” population of 10,000 diploid individuals. For each individual, we simulated a 1Mb “gene” which can accumulate eQTL mutations, along with a disjoint 100kb neutral segment that can accumulate neutral mutations at the same rate as the “gene”. Effectively, these mimic the structure of the PrediXcan models we use to characterize regulatory variants in the real data. Expression of the simulated gene is under stabilizing selection, and eQTL mutation effect sizes are drawn from a standard normal distribution. Relative fitness of individuals is calculated from total eQTL mutation effect sizes. The ancestral population is allowed to reproduce for 20,000 generations, with a mutation rate of 8 × 10^-9^ and a recombination rate of 1 × 10^-7^. The fitness optimum is centered at mean 0, standard deviation 1.

After 20k generations, we split the ancestral population into 5 subpopulations of 10,000 individuals each (P1, P2, P3, P4, P5) and the simulations run for another 400 generations. After the ancestral population split, the mutation and recombination rates are lowered to 1 × 10^-10^ and 1 × 10^-8^, respectively, and directional selection is applied to P5 by shifting the fitness optimum by a varying amount, while holding it constant for the other 4 populations. At the end of the simulations, we output the position, frequency, and effect size of the generated eQTL mutations and mutations from the neutral DNA segment. We ran 20k simulations (where each simulation represents 1 “gene”) for each of the 8 fitness optimum shift (FOS) conditions for P5 (FOS = 0.01, 0.02, 0.05, 0.1, 0.2, 0.5, 1). For each simulation, we then calculated the *Q_X_* statistic and P-value using either the gamma correction or effect permutation (drawing effect sizes from either the neutral background or the other simulations in the same FOS). We set the significance threshold to *P* < 10^-4^.

### Population Expression comparison

We predicted expression by applying the best JTI Models to the same individuals used to calculate *Q_X_* (2504 unrelated individuals from 1kG, and 929 from HGDP). We summarized across populations by taking the median predicted expression within each population.

We also compared this predicted expression observed expression in LCLs for both datasets. For 1kG, this was the same data described in *Regulatory Variant Selection*. For HGDP, this included 45 individuals from 5 geographic regions– Africa, the Middle East, East Asia, South Asia, and the Americas. We calculated the significance of the agreement across genes in the analysis as following: For each gene, we calculated a Spearman correlation between the median observed and predicted expression across all populations. We then summed the correlations together. We calculated an empirical P-value for each plot by shuffling the medians 10,000 times and repeating the summation.

### Other Selection Scores

We obtained loss-of-function (LoF) tolerance scores for each gene from Gnomad v2 (Lek et al., 2016), and used the observed/expected number of LoF variants as a measure of conservation on the protein sequence (where a low score indicates more constraint). We downloaded the phyloP100way track from the UCSC Genome Browser, and calculated the average score for each gene across the 200kb window centered around the gene using the bigWigAverageOverBed tool. Positive phyloP scores indicate greater conservation in the region.

We calculated iHS and nSL statistics in all 26 1kG populations using SelScan 2.0.0 (Szpiech, 2021). nSL was calculated using unphased genomes (Ferrer-Admetlla et al., 2014). For iHS, we used phased genomes (Voight et al., 2006), and polarized ancestral and derived alleles based on the Chimpanzee genome. We interpolated recombination maps for our sites from 1kG maps (Spence and Song, 2022). For both we focused on variants with MAF > 0.05 in the population in question, then took the mean statistic across the 200kb window around each gene.

### Enrichment Tests

We tested for enrichment for particular gene sets by calculating a ‘tiered’ enrichment. We sorted the genes by p-value, then tested for enrichment in the top N (for N = 10, 20, 30, 50, 75, 100, 150, 200, 300, 500, 1000, 2000, 3000, 5000 and 10000). Enrichment is calculated as the proportion of genes in the top N divided by the overall proportion that are in the gene set in question. We used the Binomial test to calculate a *P*-value, and calculated a confidence interval for each N using the Agresti-Coull method (Agresti and Coull, 1998). We did this for 3788 housekeeping genes (Eisenberg and Levanon, 2013) and for 2899 LoF-intolerant genes, where a gene is LoF-intolerant if the upper bound of the 95% confidence interval of the observed/expected ratio is lower than 0.35 (Lek et al., 2016), as well as 5352 genes that have undergone balancing selection on expression across species (Chen et al., 2018). We additionally tested sets of genes based on their function. These included 20 diet genes with previously-identified signals of selection (Rees et al., 2020) as well as two sets of immune genes: 1257 virus-interacting proteins (Enard et al., 2016) and 128 interferon response genes (all products in the gene ontology term GO:0032606 and all child terms).

## Data and Code Availability

*Q_X_* scores and *P*-values are available as supplementary files with the manuscript, and all other data are previously published and publicly available. Scripts for parsing data files and running analyses are available via Github.

## Acknowledgements

We thank Jeremy Berg, Ziyue Gao, ML Benton and Sarah Fong for helpful discussions, Christopher Adams for help with the 1000 Genomes data, and Lin Poyraz for a script to polarize iHS scores.

## Author Contributions

L.L.C. and I.M. designed the experiments and wrote the manuscript. F.C.R-A. did the simulations and power analyses. L.L.C conducted all other experiments and analyzed data. All authors edited and approved the manuscript.

## Supplementary Materials

**Supplementary Figure 1:**
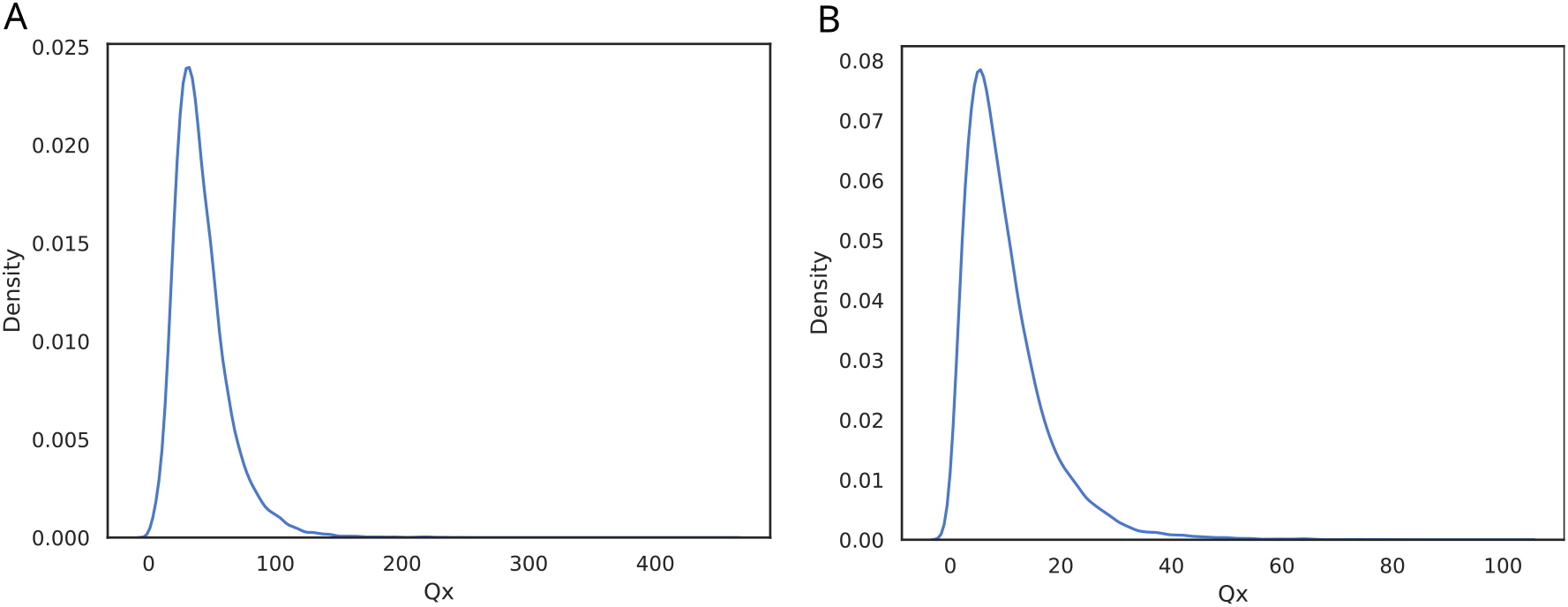
Distributions of *Q_X_* scores for the Best Models of all protein-coding genes in A) 26 1kG B) 7 HGDP populations.

**Supplementary Figure 2:**
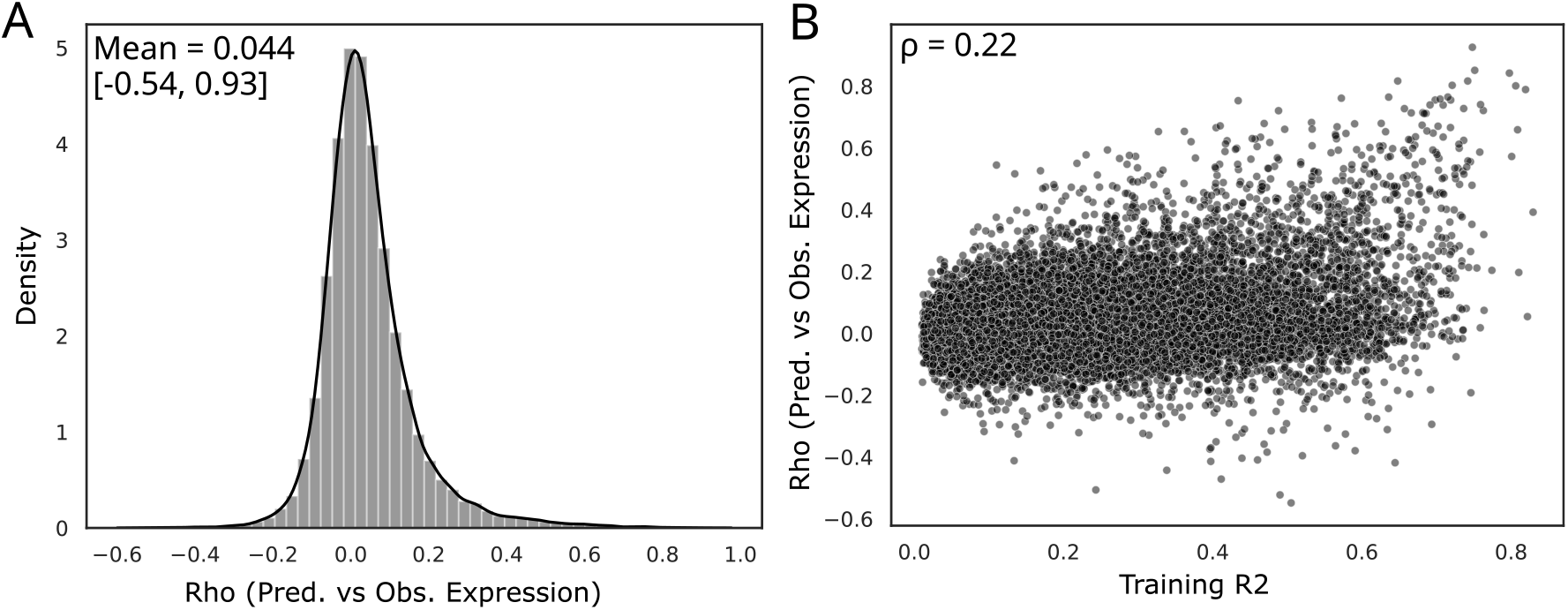
A) Rank correlation between observed and predicted expression for the Best Models of all protein-coding genes in LCLs B) Scatterplot of that rank correlation vs *R*^2^ from the JTI training process.

**Supplementary Figure 3:**
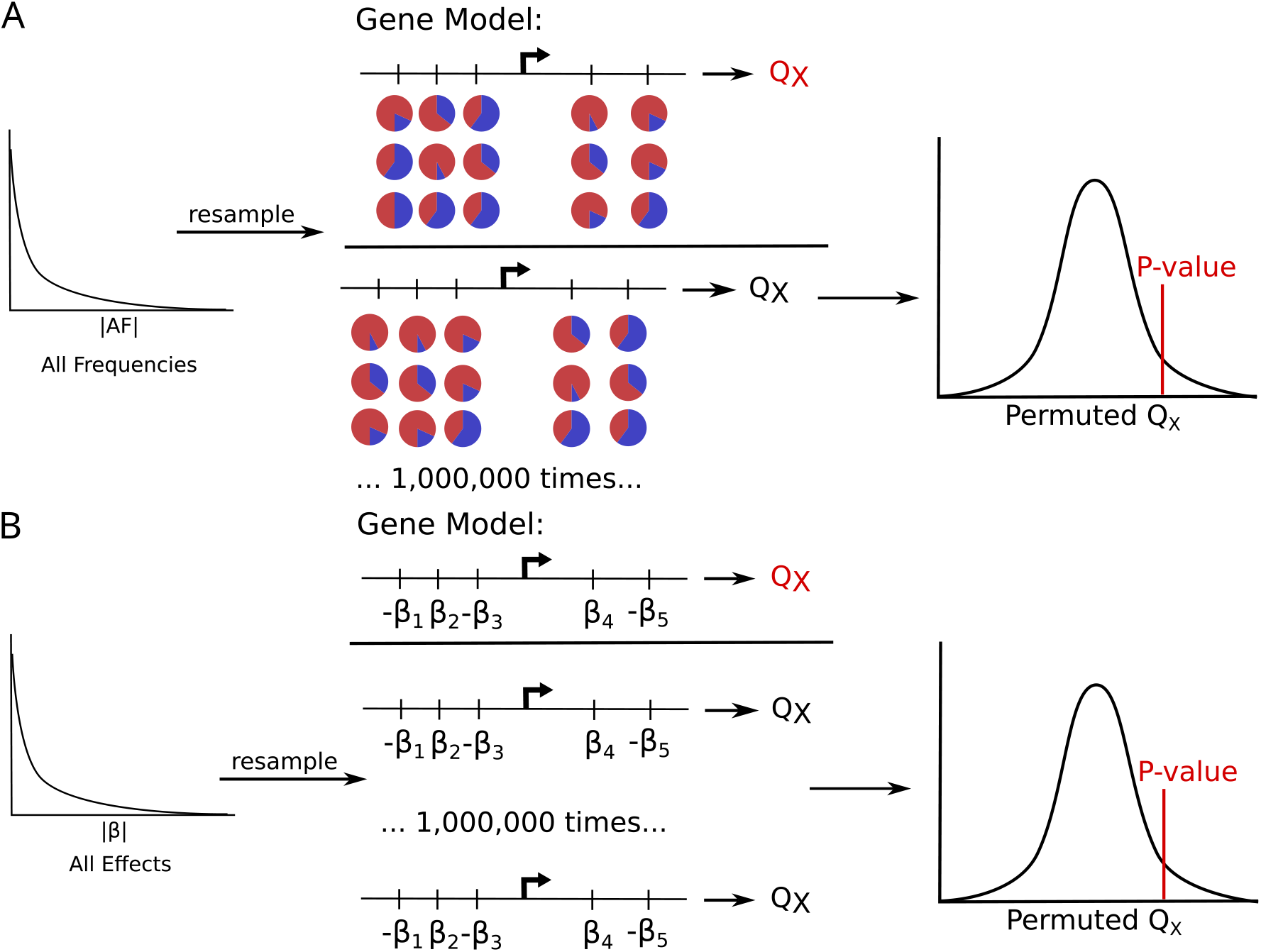
Schematics describing the two permutations schemes we tried. A) One option is to calculate an empirical *P*-value by permuting the variant allele frequencies across a model. B) The other option is to instead permute the effect sizes of the variants, while holding effect direction constant. In both, we resampled 100,000 times for each gene, extending to 1,000,000 for those with *P* < 10^-4^.

**Supplementary Figure 4:**
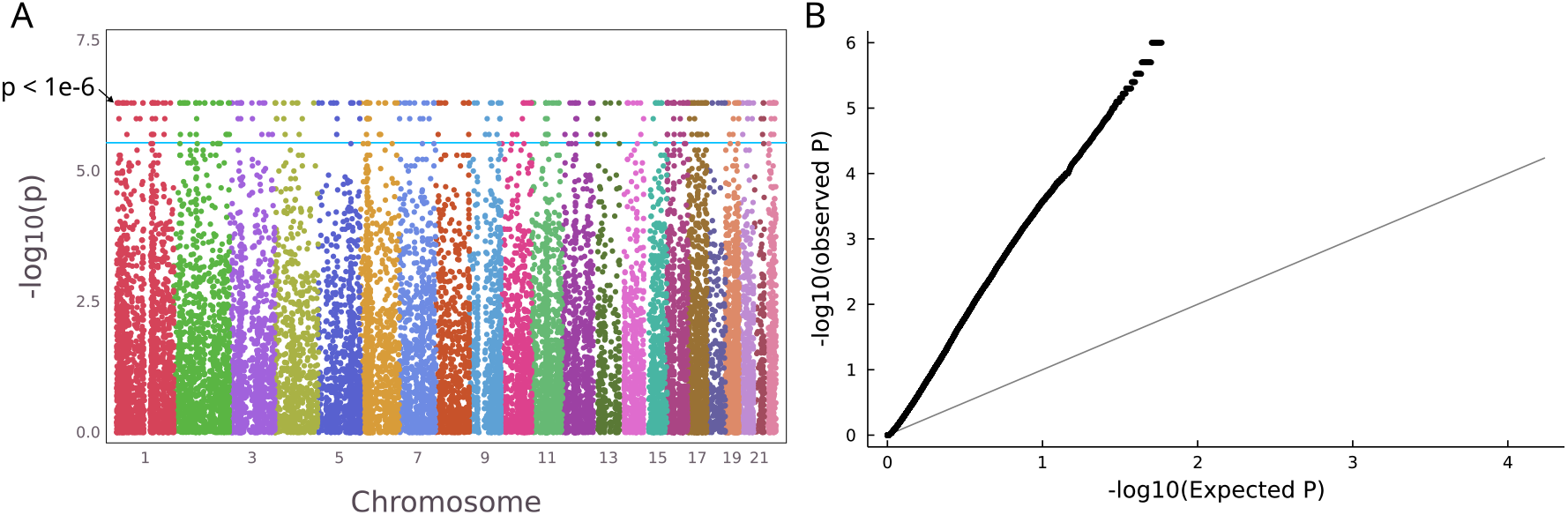
P-values for the Best Model protein-coding genes calculated by permuting the allele frequencies of the model variants. A) Manhattan plot; the maximum of *−log*_10_(*P*) is where *P* < 1,000,000. B) a QQ plot of the P-values, excluding those for which *P* < 1,000,000.

**Supplementary Figure 5:**
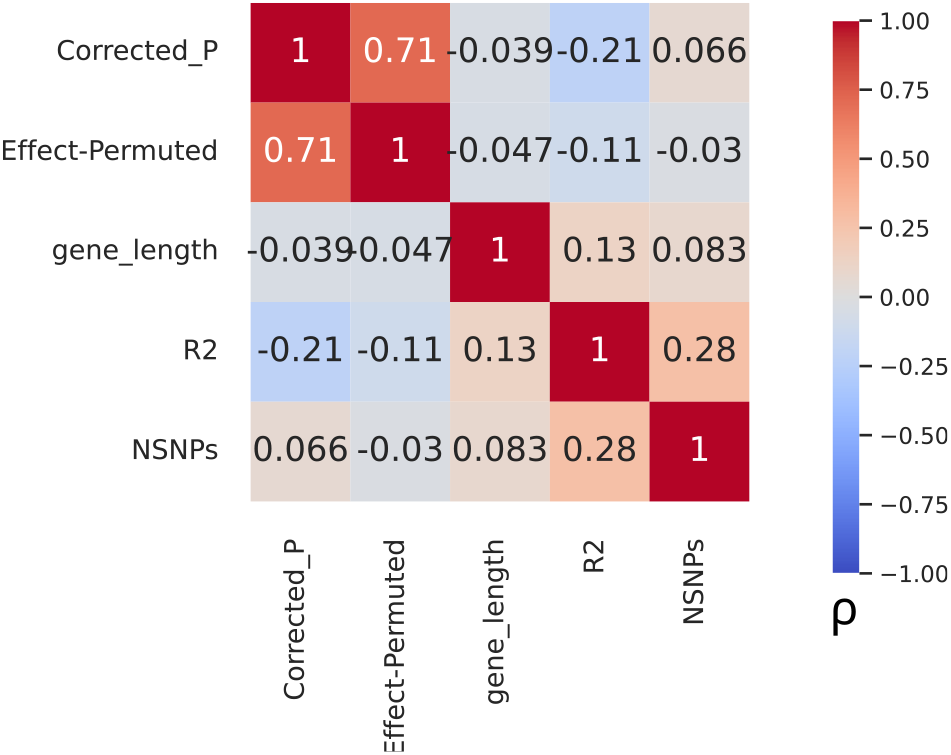
*Q_X_* is correlated with technical aspects of JTI models. Pairwise Spearman rank correlations between gamma-corrected P-values and effect size permutation P-values with various technical variables. *R*^2^ and *NSNPs* refer to the training performance of the Best JTI models used to calculate *Q_X_*

**Supplementary Figure 6:**
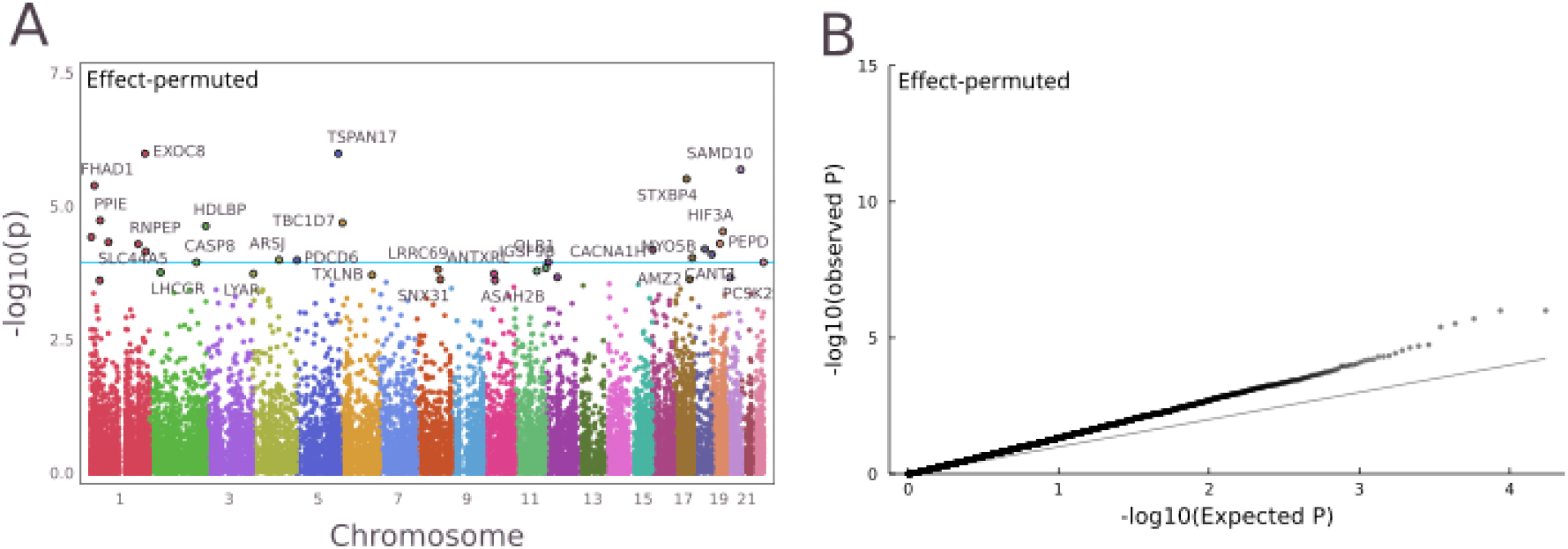
Effect-Permuted P-values. A) Manhattan and B) QQ plots for effect size permuted *P*-values (Methods). The blue line in each Manhattan plot corresponds to the threshold where FDR < 0.1.

**Supplementary Figure 7:**
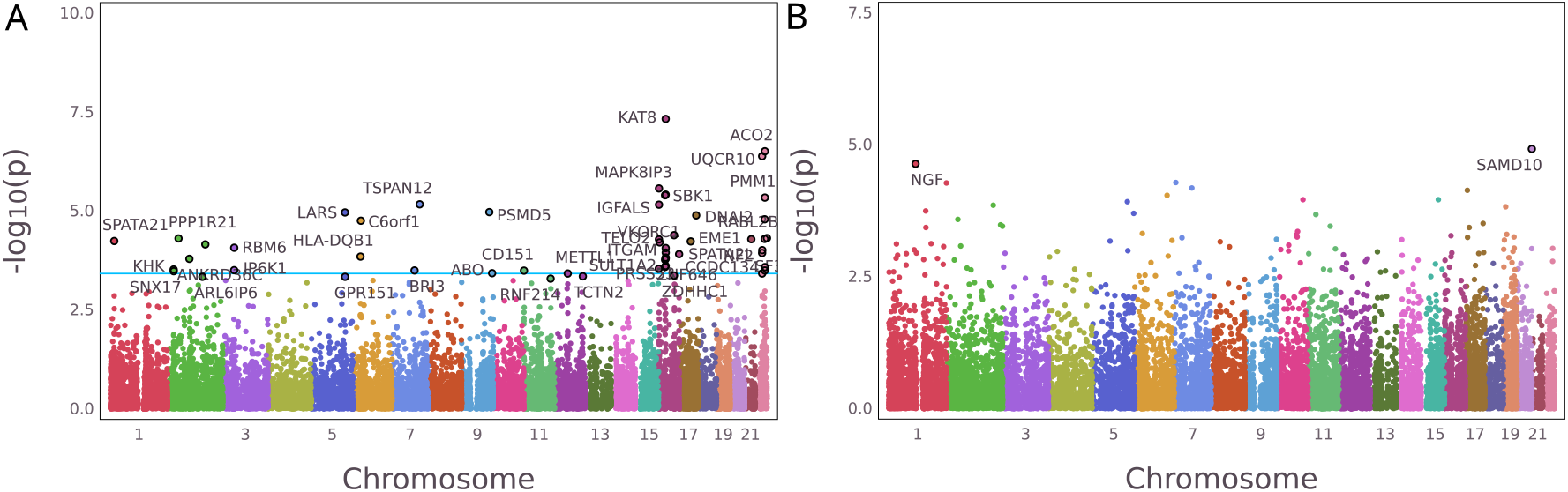
HGDP analysis. Manhattan plots of A) gamma-corrected *P*-values and B) effect size permuted *P*-values for *Q_X_* calculated in HGDP. Blue line corresponds to FDR < 0.1.

**Supplementary Figure 8:**
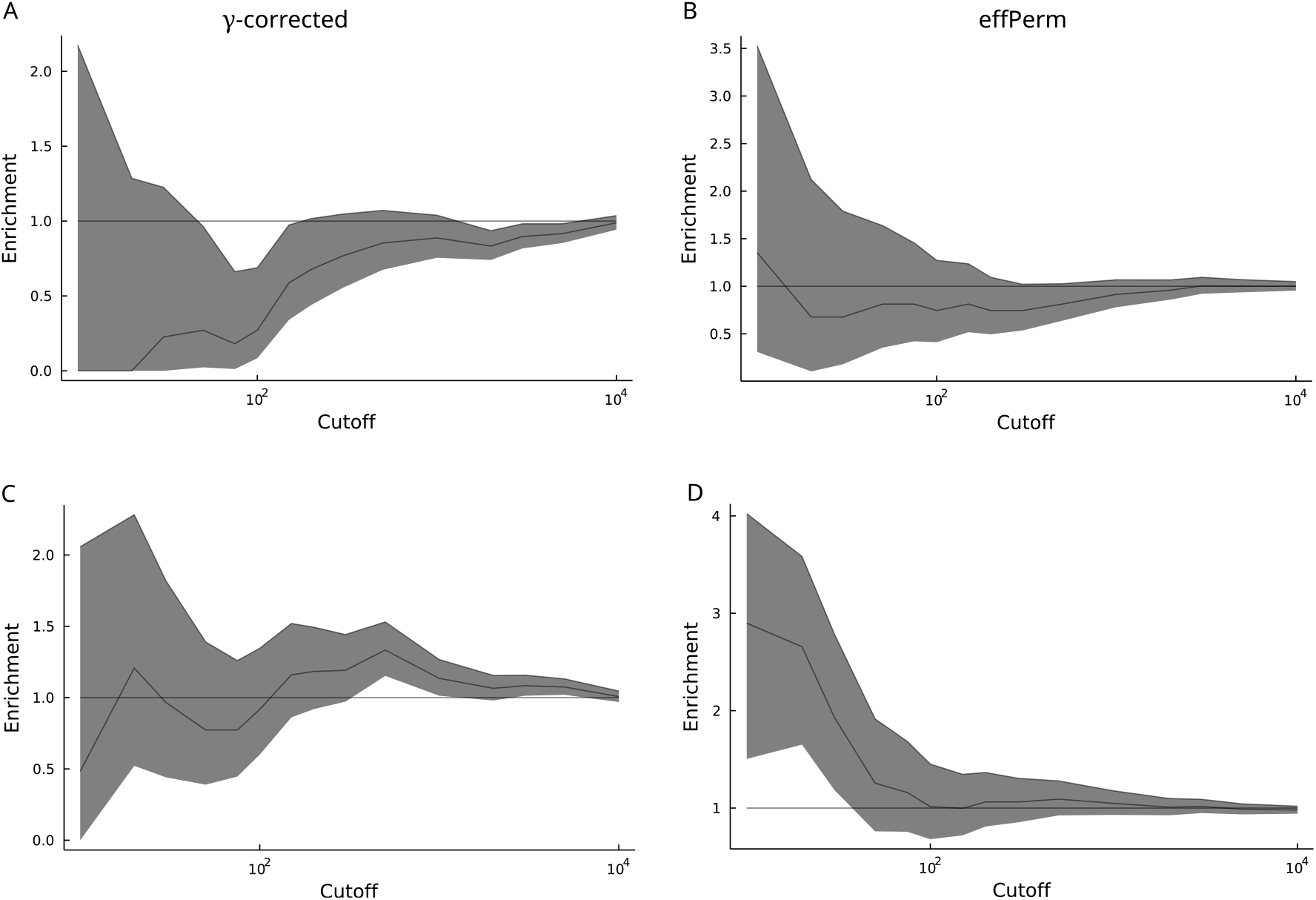
Predicted regulation for genes with *FDR* < 0.05 using gamma-corrected and effect size permuted P-values. A) and B) LoF intolerant, C) and D) Housekeeping genes

**Supplementary Figure 9:**
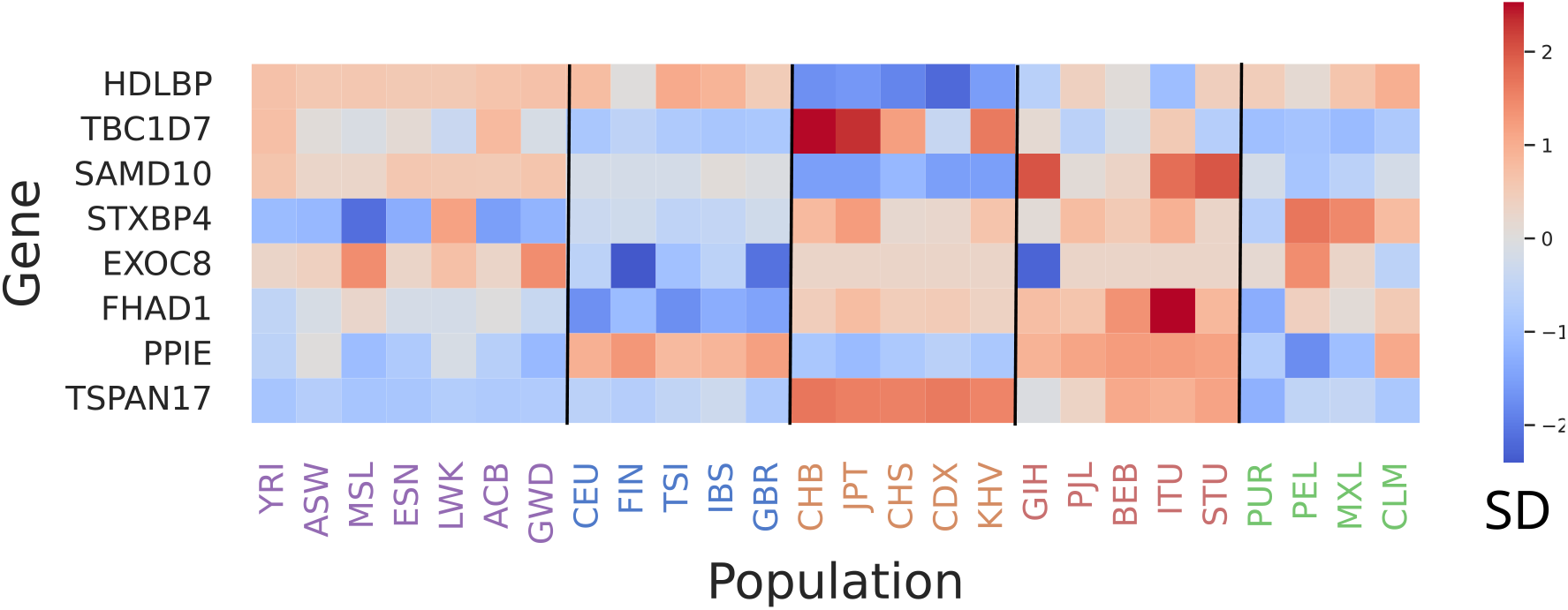
Predicted expression for genes with *FDR* < 0.05 using effect size permuted P-values. Each square is coloured by the median predicted expression for that gene in that population, and is standardized across the row.

**Supplementary Figure 10:**
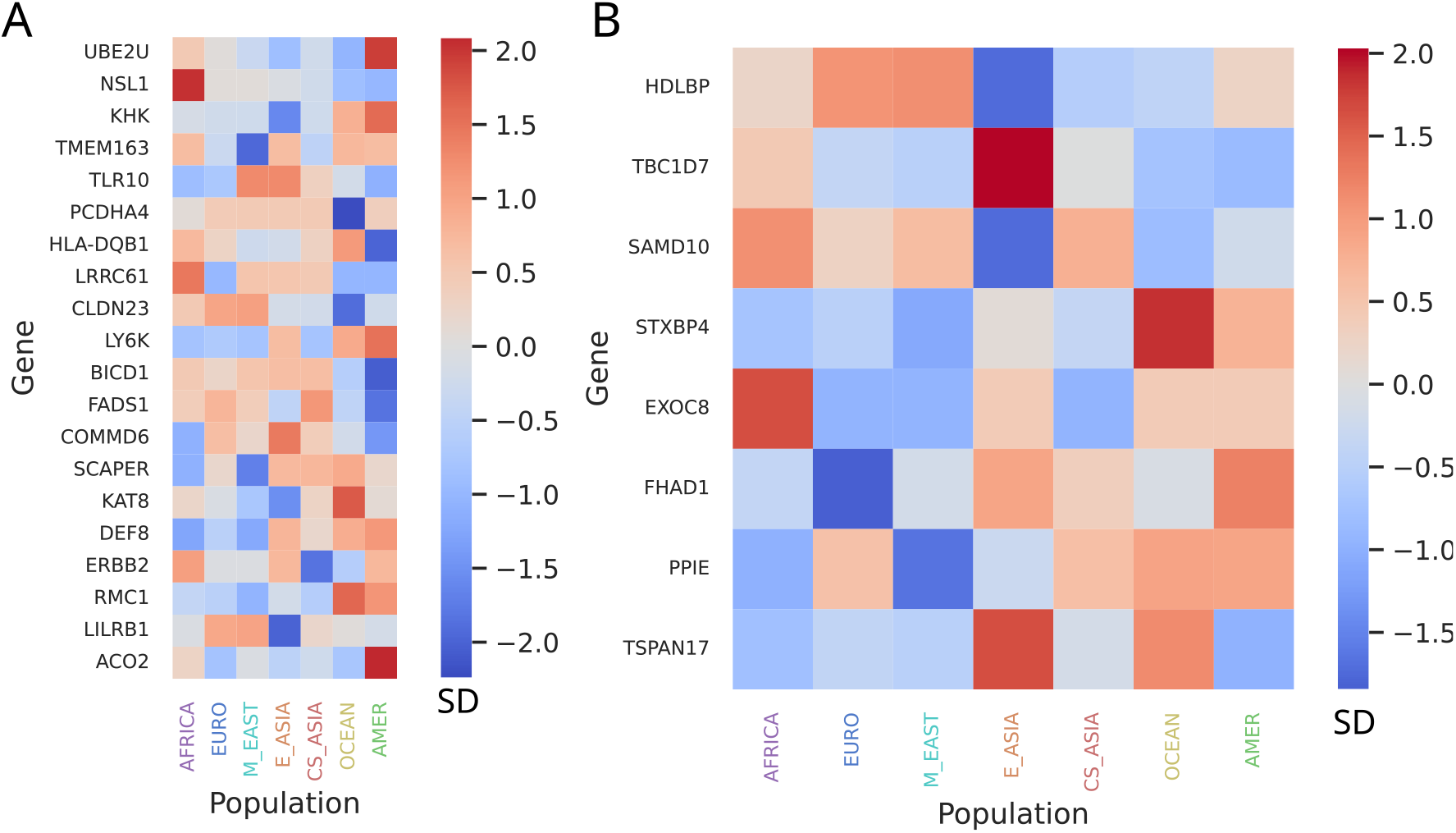
HGDP Predicted expression patterns largely agree with 1kG across populations. Heatmap of median predicted expression in each HGDP population A) the top genes in each ‘peak’ based on the gamma-corrected *P*-values, and B) genes with *FDR* < 0.05 for the effPerm *P*-values in 1kG. For display purposes, values are standardized across populations for each gene.

**Supplementary Figure 11:**
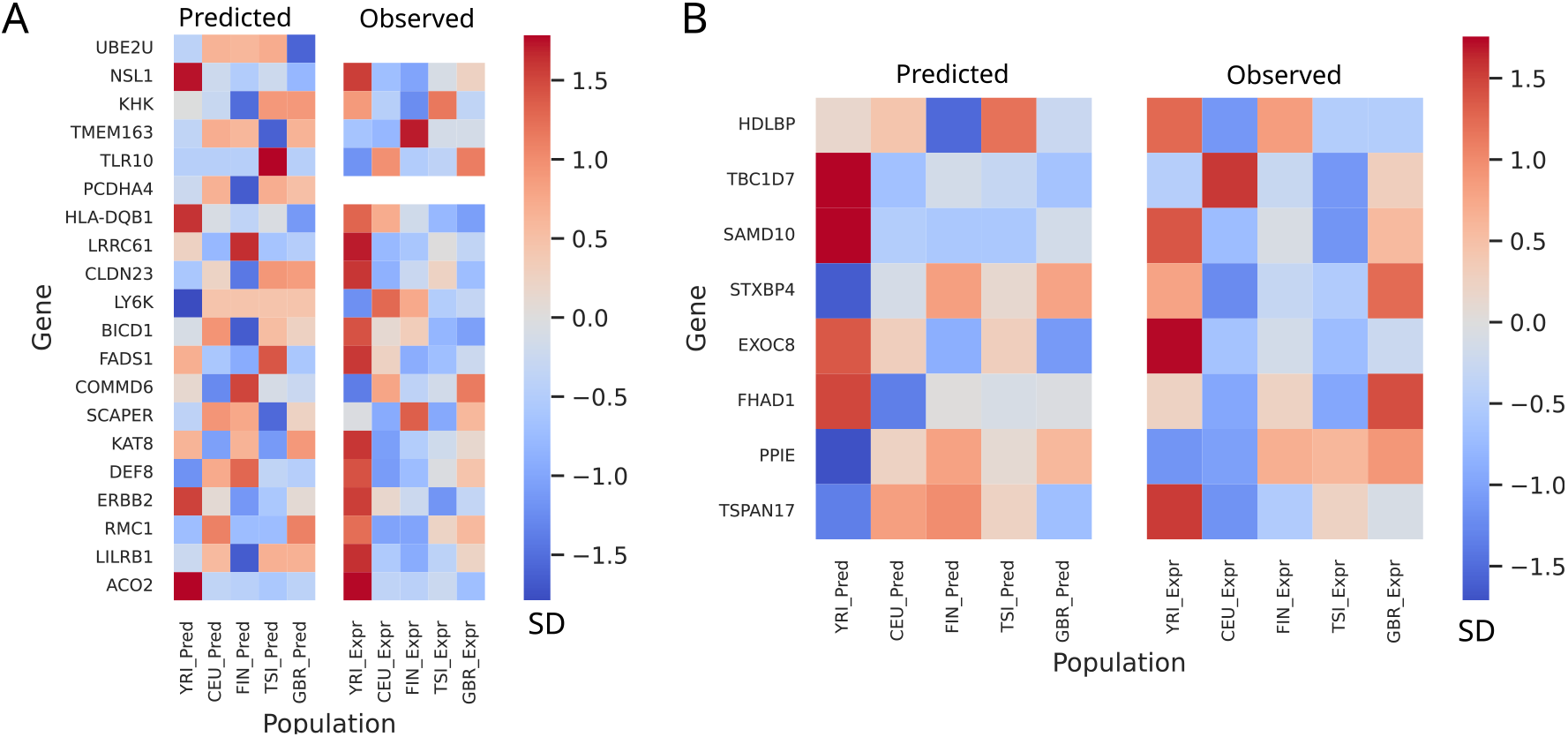
Predicted expression pattern does not necessarily correlate with observed expression in LCLs. Heatmaps of median predicted and observed expression in each 1kG population for A) the top genes in each ‘peak’ based on the gamma-corrected *P*-values (*P* = 0.241), and B) genes with *FDR* < 0.05 for the effPerm *P*-values (*P* = 0.717). For display purposes, values are standardized across populations for each gene.

**Supplementary Figure 12:**
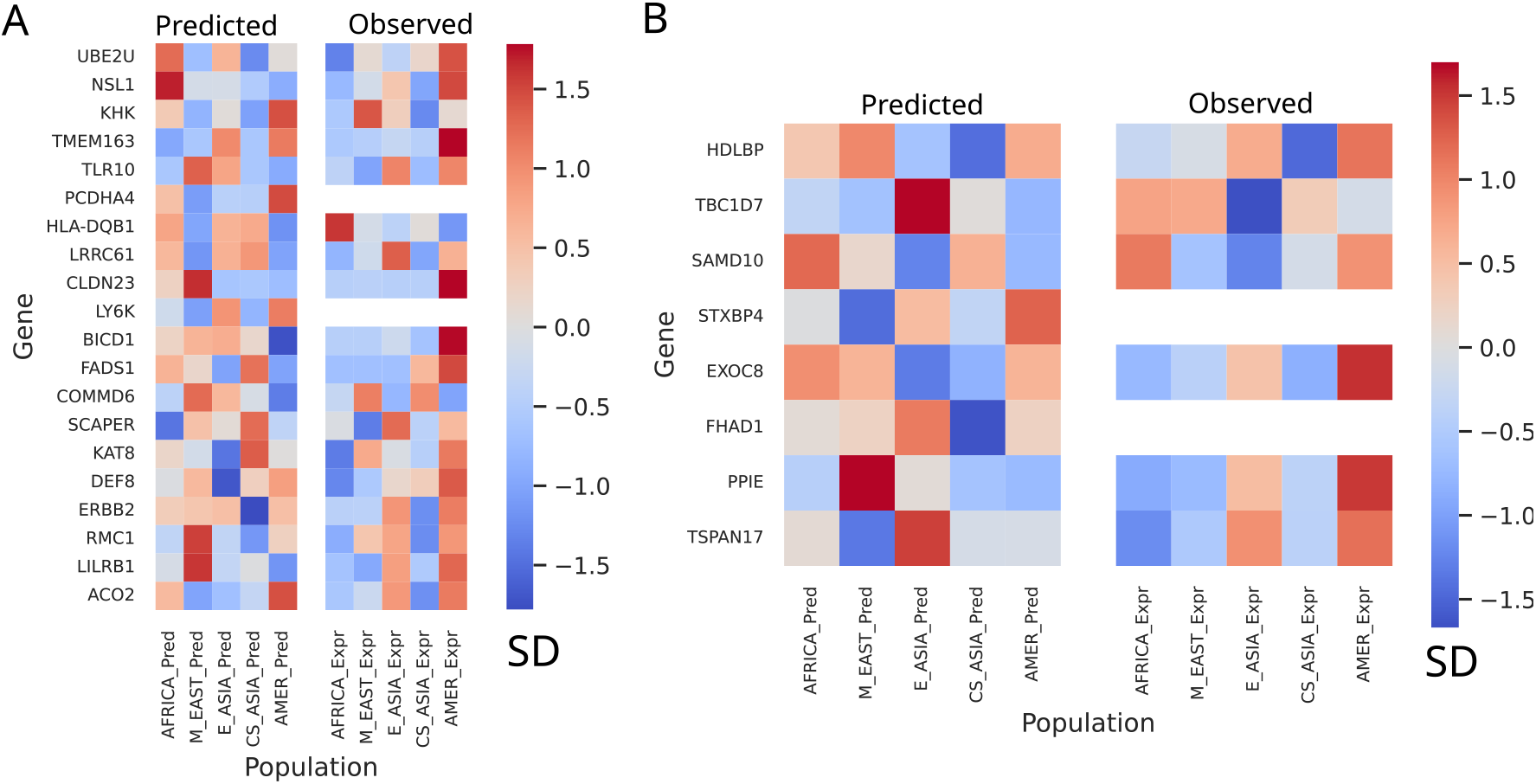
Predicted expression pattern does not necessarily correlate with observed expression in LCLs. Heatmaps of median predicted and observed expression in each HGDP population for A) the top genes in each ‘peak’ based on the gamma-corrected *P*-values (*P* = 0.612), and B) genes with *FDR* < 0.05 for the effPerm *P*-values (*P* = 0.697). For display purposes, values are standardized across populations for each gene.

**Supplementary Figure 13:**
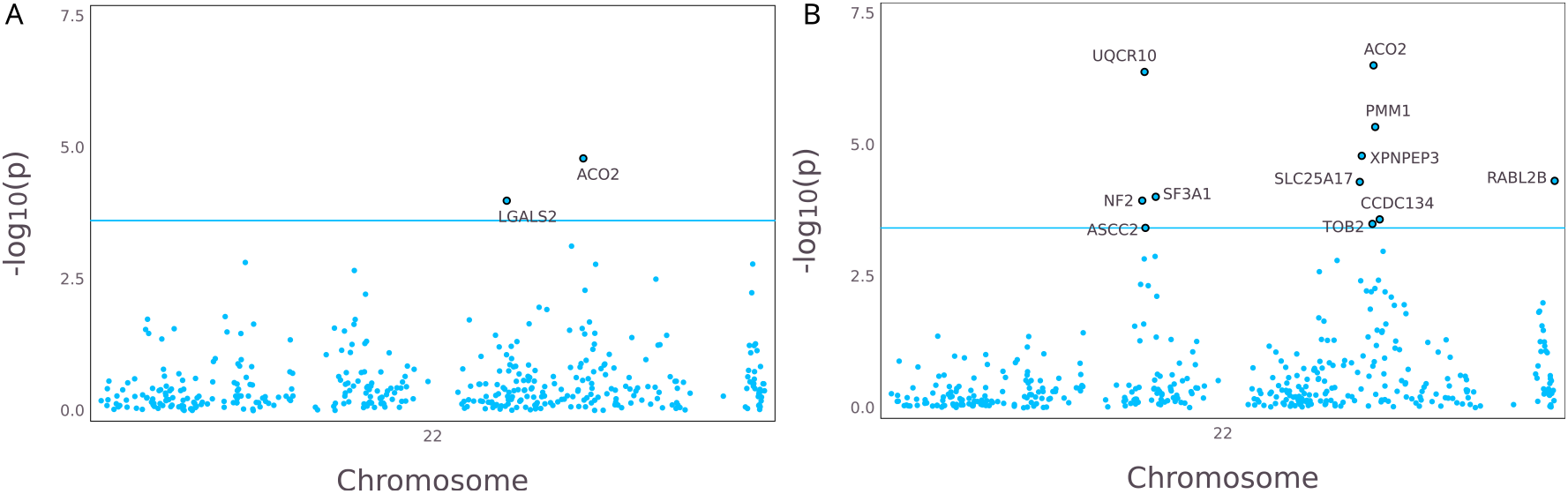
*ACO2* is under selection in 1kG and HGDP Zoomed Manhattan plots of chr22 for A) 1kG and B) HGDP.

**Supplementary Figure 14:**
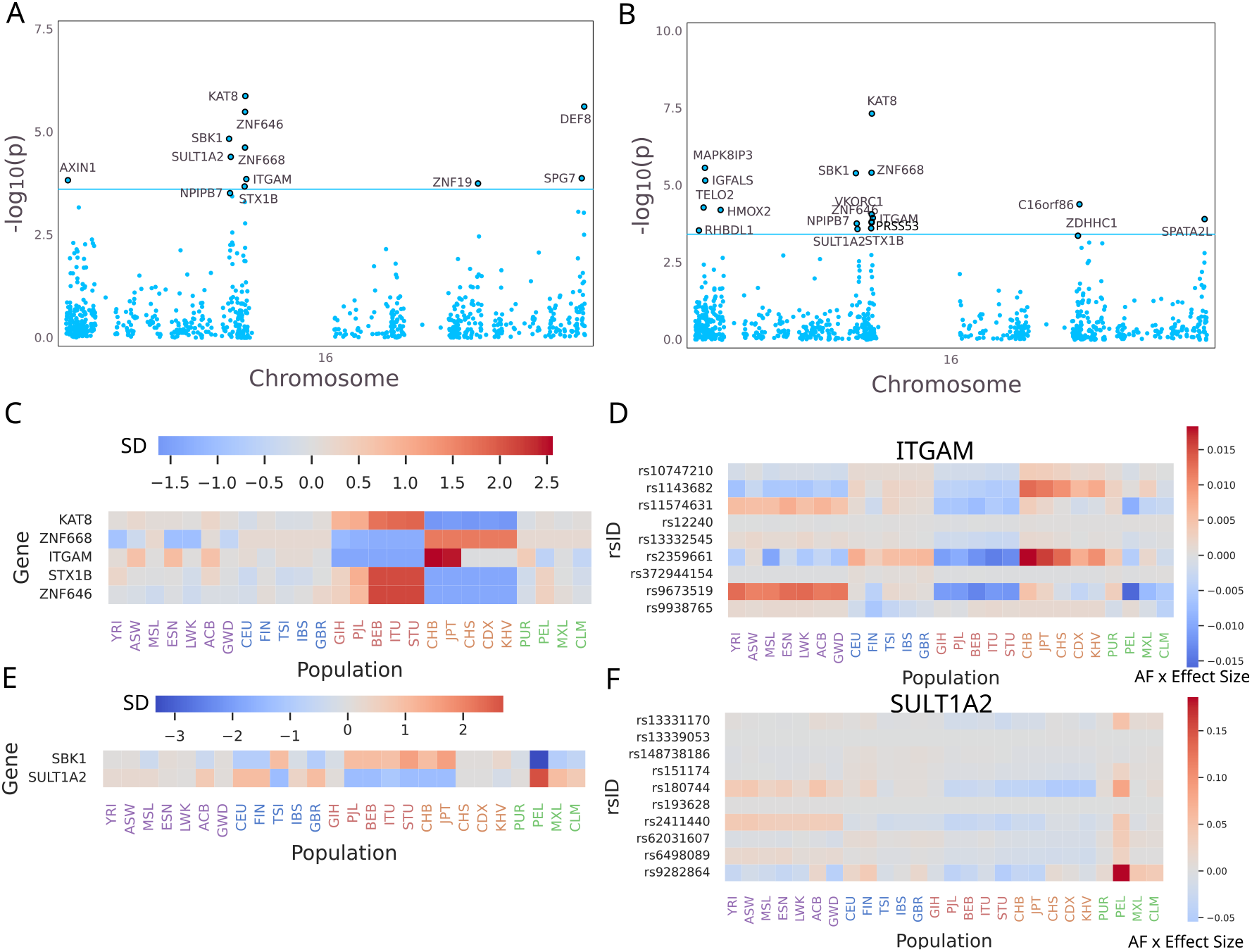
the *KAT8* peak is actually 2 independent signals. Zoomed Manhattan plots of chr16 for A) 1kG and B) HGDP. C) Median Predicted expression for the 5 replicating genes near *KAT8*; D) Heatmap of the product of JTI effect size times effect allele frequency for variants in the *ITGAM* model. E) Median predicted expression for SBK1 and SULT1A2, which replicated in 1kG and HGDP, and F) Heatmap of the product of JTI effect size times effect allele frequency for variants in the *SULT1A2* model.

